# Cholinergic-estrogen interaction underpins the effect of education on attenuating cognitive sex differences in a Thai healthy population

**DOI:** 10.1101/2022.11.10.515954

**Authors:** Chen Chen, Bupachad Khanthiyong, Sawanya Charoenlappanit, Sittiruk Roytrakul, Samur Thanoi, Sutisa Nudmamud-Thanoi

## Abstract

The development of human brain is shaped by both genetic and environmental factors. Sex differences in cognitive function have been found in humans as a result of sexual dimorphism in neural information transmission. Numerous studies have reported the positive effects of education on cognitive functions. However, little work has investigated the effect of education on attenuating cognitive sex differences and the neural mechanisms behind it based on healthy population. In this study, the Wisconsin Card Sorting Test (WCST) was employed to examine sex differences in cognitive function in 135 Thai healthy subjects, and label-free proteomic method and bioinformatic analysis were used to study sex-specific neurotransmission-related protein expression profiles. The results showed a sex difference in two WCST subscores: percentage of Total corrects and Total errors in the primary education group (Bayes factor>100) with males performed better, while such differences eliminated in secondary and tertiary education level. Moreover, 11 differentially expressed proteins (DEPs) between men and women (FDR<0.1) were presented in both education groups, with majority of them upregulated in females. Half of those DEPs interacted directly with nAChR3, whereas the other DEPs were indirectly connected to the cholinergic pathways through interaction with estrogen. These findings implied that Cholinergic-estrogen interaction underpins the effect of education on attenuating cognitive sex differences in a Thai healthy population.

## 1. Introduction

Human brain development is influenced by both genetic and environmental factors. The X and Y chromosomes define not just a person’s sex, but also the morphological and functional distinctions between male and female brains. For example, the male brain has more volume and a higher proportion of white matter (1), but the female brain has a bigger corpus callosum (2). According to Ingalhalikar et.al. (3), the male brain is thought to be intrahemispherically connected, whereas the female brain is optimized for interhemispheric connectivity. There is growing acknowledgment that the human brain is sensitive to environmental circumstances throughout development, such as socioeconomic status, education, nutrition, and parental care (4). Evidence suggests that children from higher-income families have greater volumes of grey and white matter in the inferior frontal gyrus and bigger hippocampus sizes than their lower socioeconomic status peers (5, 6). Furthermore, education not only enhances human brain development but also attenuates cognitive deficits that occur with aging (7-9). Schooling increases cortical grey matter volumes in several brain areas (10), and even a few years of education has been demonstrated to contribute to brain cognitive reserve (11).

Under the joint actions of genetic and environmental factors, male and female brains may differ in terms of cognitive strategies and/or cognitive styles (12). These include different learning style preferences (13), as well as sex differences in decision-making styles (14).

There has been minimal research into the effect of education on attenuating cognitive sex differences. Bloomberg, Dugravot (15) reported in sex comparisons in adults in England, males in the low education group and the earliest birth cohort performed better in a semantic fluency test, whereas females in the higher education group and the latest birth cohort achieved higher fluency scores in the same test. This finding implies that schooling, as well as secular changes in education level across birth cohorts, play a role in determining cognitive performance in females. However, given this result was based on data collected from participants at the age of 60 years, it is unknown whether such a result extends to other age groups and other domains of cognition.

Cognitive performance is the reflection of information transmitted along with neural circuits of brain areas (16, 17), and learning and obtained experiences have been documented to alter the morphology of synapses as well as promote their plasticity (18). The study of how education affects neurotransmission in male and female brains is still in its infancy and deserves more attention. Therefore, the purpose of this study is to investigate the effects of education on cognitive sex differences in a healthy Thai population with a wider range of birth cohorts, as well as to investigate the neural mechanisms behind such effects.

## 2. Materials and methods

### 2.1 Study subjects

As described in a previous study from our lab (19), one hundred and thirty-five healthy volunteers between the ages of 22 and 70 were recruited. Subjects were divided into two categories based on their education level: those who had acquired no more than primary education, those who had received secondary education and, for some, tertiary education. Subjects with abnormal mental health evaluated by the Thai Mental Health Indicator (THMI-55) were excluded from the current study. The Mini-Mental State Examination (MMSE) was also used to screen out participants with dementia. To limit the likelihood of confounding by population stratification, all individuals were Thai ethnicity (20). All participants signed informed consent forms. The experimental protocols were approved by the Human Ethics Committee of Naresuan University (COA No. 0262/2022).

### 2.2 Cognitive assessment

The Wisconsin Card Sorting Test (WCST) has long been used in clinical and research settings to assess frontal cortex (FC) function (21). The recent neuroimaging study revealed that, in addition to FC, a widespread network of prefrontal, frontal, temporal, frontal-temporal, parieto-occipital regions are activated during WCST (22), and multiple domains of cognitive function including but not limited to working memory, attention, cognitive set-shifting, planning or organizing, problem-solving and decision-making are activated during the process of WCST (23).

In this study, all subjects were tested using a computer-based WCST (Inquisit 3.0.6.0), and the raw score was analyzed to reflect different domains of cognitive function, as shown below (24):

- The percentage of total corrects (%Corrects): the entire number of correct response cards multiplied by 100 and divided by the total number of cards, reflecting initial conceptualization and attention.
- The percentage of total errors (%Errors): the total number of incorrect response cards multiplied by 100 and divided by the entire number of cards, reflecting nonspecific cognitive impairment.
- The number of categories completed (Category completed): determined by applying the score range from 1 to 6, reflecting cognitive set-shifting.
- The perseverative errors (PE): the score was used to assess the inability to correct a response due to ignorance of relevant stimuli, reflecting cognitive inflexibility.
- Trails to complete the first category (1^st^ Category): the score ranges from 0 to 128, which is the number required to complete the initial category of the task.

### 2.3 Blood sample collection

A 3ml cubital vein blood was collected from each enrolled subject in a clotted blood tube, and the blood was centrifuged at 3,000 rpm for 5 minutes. The serum was then transferred into a 1.5ml microcentrifuge tube and stored at -80°C in a refrigerator for future use, all samples were coded to ensure anonymity.

### 2.4 Label-free quantitative proteomics analysis

The label-free quantitative proteomics analysis was used to compare serum protein expression profiles between males and females. The analytic processes were performed by the Proteomics unit from National Centre for Genetic Engineering and Biotechnology, Pathum Thani, Thailand, which include protein digestion, Liquid Chromatography with tandem mass spectrometry (LC-MS/MS) analysis, protein identification, and protein quantitation.

In brief, the protein concentration of all serum samples was determined using the Lowry assay with BSA as a standard protein (25). Five micrograms of protein samples were digested with sequencing grade porcine trypsin (1:20 ratio) for 16 hours at 37 °C. The tryptic peptides were dried in a speed vacuum concentrator and resuspended in 0.1% formic before further analysis.

The prepared tryptic peptide sample of each subject was injected individually into an Ultimate3000 Nano/Capillary LC System (Thermo Scientific, UK) coupled to a Hybrid quadrupole Q-Tof impact II™ (Bruker Daltonics) equipped with a Nano-captive spray ion source. Mass spectra (MS) and MS/MS spectra were obtained in the positive-ion mode at 2 Hz over the range (m/z) 150–2200. To minimize the effect of experimental variation, three independent MS/MS runs were performed for each sample.

MaxQuant 1.6.6.0 was used to quantify and identify the proteins in each sample with the Andromeda search engine to correlate MS/MS spectra to the Uniprot *Homo sapiens* database (26). The proteins were identified using a 10% protein false discovery rate (FDR), carbamidomethylation of cystein as fixed modification, and the oxidation of methionine and acetylation of the protein N-terminus as variable modifications. Only proteins with at least two peptides, and at least one unique peptide, were considered as being identified and used for further data processing.

### 2.5 Bioinformatic analysis

Before any analysis, data cleansing and preprocessing were performed by using Perseus ver. 1.6.15.0 (27). A list containing all proteins identified previously was submitted to jvenn (web application) to explore those proteins shared by males and females at both primary and secondary and tertiary education groups (28). Differentially expressed proteins (DEPs) between males and females in each education group were detected independently using the Linear Model for Microarray Data (LIMMA) approach within R-programming ver. 4.1.2 (29), with the FDR set at 10%. Multi-Experiment Viewer (MeV, ver.4.9.0) software was employed to illustrate the expressions of shared DEPs in men and women from both education groups (30). Protein-protein interaction (PPI) network, as well as the relationship between DEPs and cognitive function, was exhibited by Pathway Studio ver. 12.5 (31).

### 2.6 Statistical analysis

Using R-programming ver. 4.1.2, a General Linear Model (GLM) approach paired with Bayesian statistics was applied to analyze differences in WCST scores between men and women at each education level (32), with age as a covariate. Bayes factor was offered to illustrate the likelihood of supporting the alternative hypothesis (33).

## 3. Results

### 3.1 Demographic data

The Subjects were 70 males and 65 females, with a mean age of 57.75±10.35 years (range, 22-70 years). Table 1 describes their demographic data, including sex, age, and educational level. The age differences between males and females in primary, secondary and tertiary education groups were supported by weak and substantial evidence, respectively. However, the age difference between the two education groups was discovered with decisive evidence (BF>100, 95%CI=[9.65, 16.0]).

**Table 1.** Demographic data of subjects

|  | Male | Female | BF | 95% CI |
| --- | --- | --- | --- | --- |
| **Primary education** |  |  |  |  |
| No. of subjects | 49 (70%) | 42 (64.6%) |  |  |
| Age | 61.4±6.02 | 62.5±6.81 | 1.56 | [-3.78, 1.66] |
| **Secondary and Tertiary education** |  |  |  |  |
| No. of subjects | 21 (30%) | 23 (35.4%) |  |  |
| Age | 51.1±13.2 | 47.3±9.75 | 5.73 | [-3.18, 10.88] |

Data was presented as mean±SD by General Linear Model.
BF=Bayes Factor 95% CI= 95% Credible interval of difference male against female

### 3.2 Effects of education on sex differences in cognitive performance

Education was found to have potential compensatory effects on three out of the five WCST sub-scores: %Corrects, %Errors, and 1^st^ Category. In the primary education group, male scored better in %Corrects (BF>100, 95%CI= [2.74, 12.25], Cohen’s d=0.68) and had lower %Errors (BF>100, 95%CI= [-12.2, -2.71], Cohen’s d=0.67) with close to large effect size (34). While sex differences in those two scores reversed in the higher education group, however, there was only weak evidence to support the alternative hypotheses (both BF<2) (see Fig.1). Regarding the score 1^st^ Category, females were found to have fewer trials to complete the first category in the WCST test compared to males in both education groups, but there was weak evidence to reject the null hypothesis in the lower education group (BF=1.02), however, such sex difference was decisive in the second and tertiary education group (BF>100, 95%CI= [-6.65, 15.8]), although the effect size was small (Cohen’s d=0.25). Furthermore, there was insufficient evidence to accept the sex differences in score PE and Category Completed in both education groups (all BF>6).

**Fig.1.**
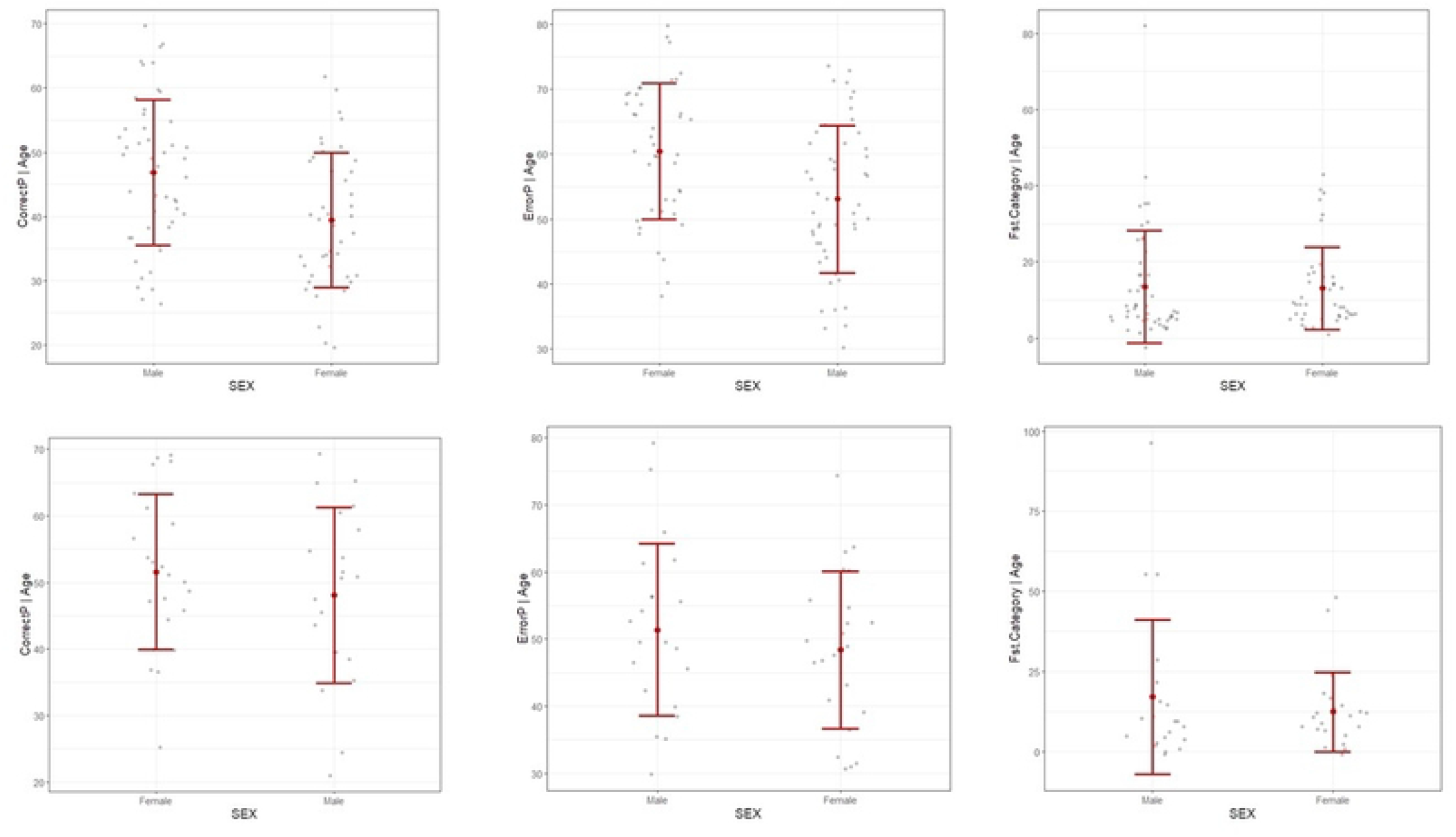
Effects of education on sex differences in WCST subscore %Corrects, %Errors, and 1^st^ Category. The upper graphs show sex differences in those scores in the primary education group, the lower graphs depict the sex differences in the same scores in the secondary and tertiary education group. CorrectP=%Corrects, ErrorP=%Errors, FstCategory=1^st^ Category.

### 3.3 Identification and relative quantification of differentially expressed proteins between men and women

After applying the criteria outlined in section 2.4, a total of 886 proteins were identified using a label-free quantitative proteomics approach, with 808 of them shared by both sexes (see Fig 2). Of those 808 proteins, we found that 11 differentially expressed proteins (DEPs) between men and women (FDR<0.1) were presented in both education groups (see Fig. 3). Except for Hematopoietic progenitor cell antigen CD34 (CD34), all of the 11 DEPs were upregulated in females.

**Fig 2:**
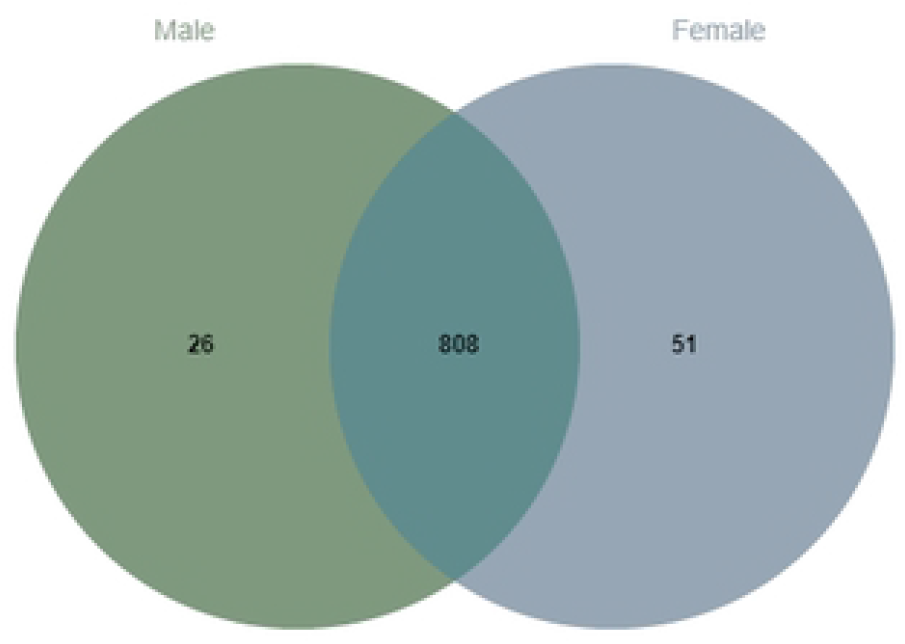
Venn diagram showing 808 proteins shared by both males and females.

**Fig. 3.**
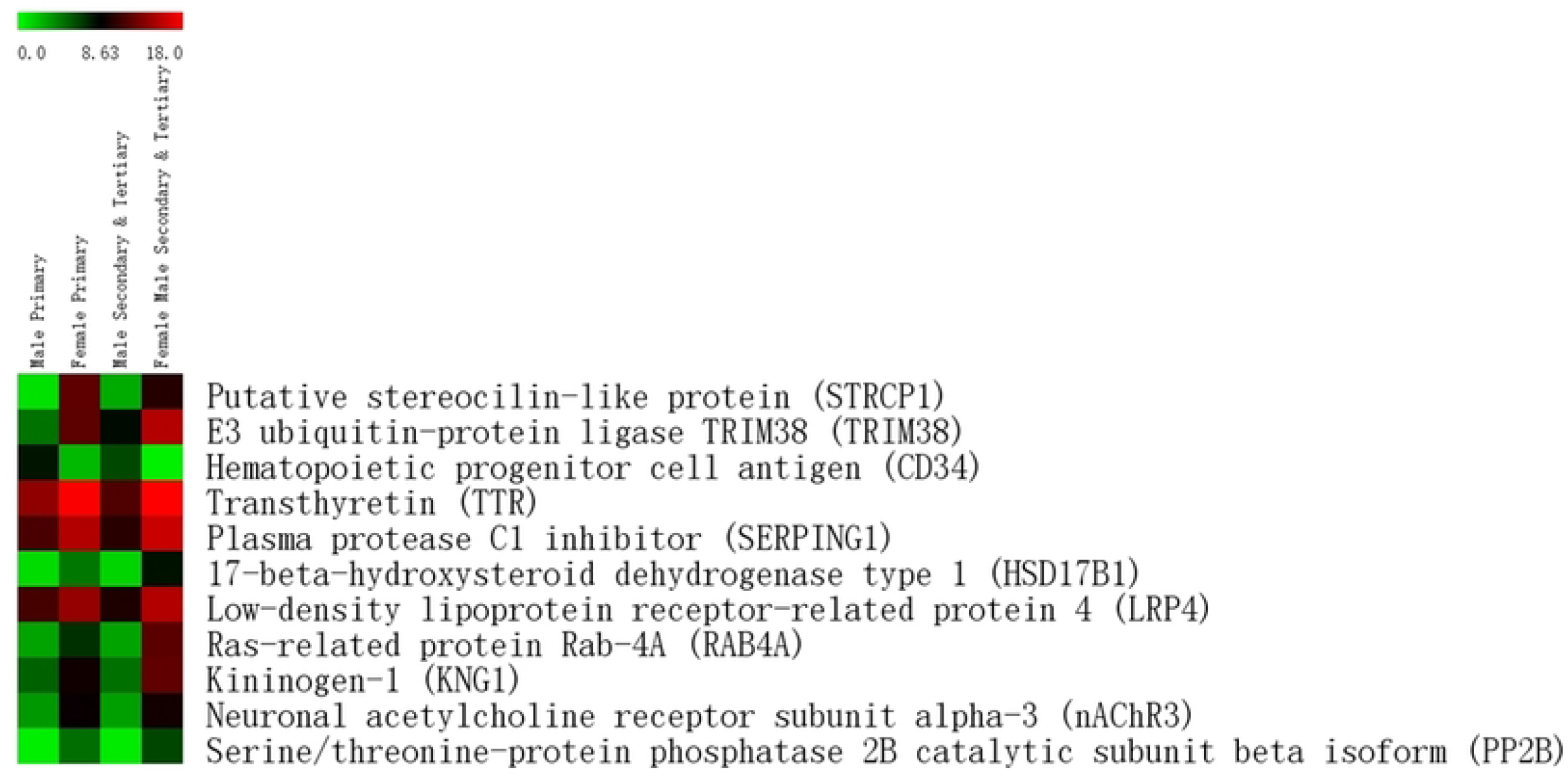
Expression heatmap of 11 proteins exhibited in both education groups.

### 3.4 Protein-protein interactions of DEPs and their relationship with cognitive function

As shown in Fig 4, out of the 11 DEPs, LRP4, TTR, TRIM38, and PP2B interacted directly with nAChR3, a subunit of nicotinic acetylcholine receptor that positively regulates cognition. While CD34, SERPING1, HSD17B1, KNG1, and TTR are indirectly connected to the cholinergic pathways through interaction with estrogen. This suggested that the cholinergic-estrogen interaction may have an influence on cognitive processes. STRCP1 and RAB4A, on the other hand, showed no interaction with other DEPs or association with cognitive function.

**Fig. 4.**
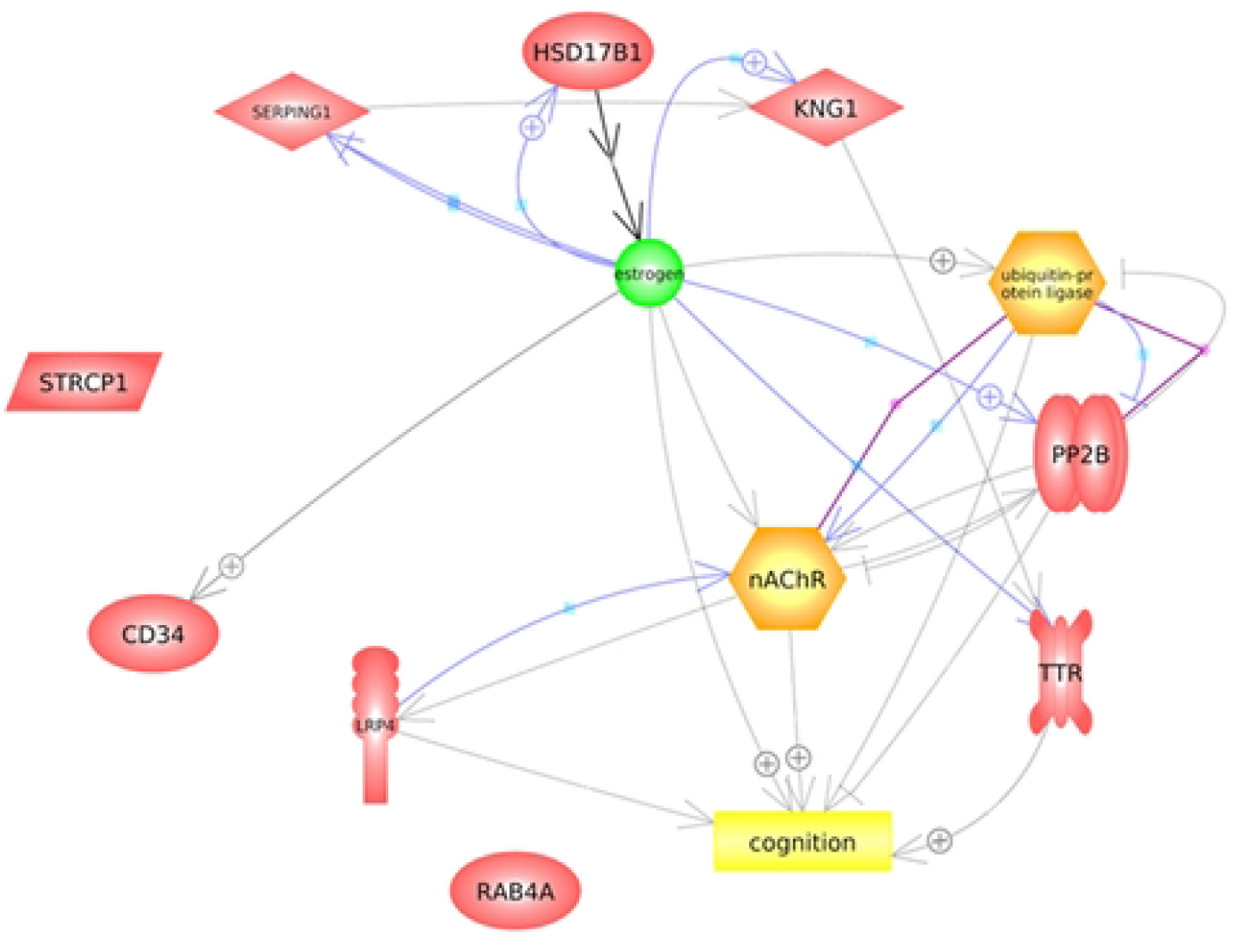
Protein-protein interaction network produced by Pathway Studio. The purple line indicates binding, the blue line indicates expression, the black line indicates chemical reactor, the solid gray indicates molecular transport, while the gray dotted line indicates regulation. The plus sign indicates positive regulation.

## 4. Discussion

In this study, we found that education may have an effect on attenuating cognitive sex differences as assessed by the WCST, with sex differences in WCST subscores %Corrects and %Errors identified in the lower education group but not supported by sufficient evidence in the higher education group. This finding is consistent with previous research indicating that sex differences in attention control to spatial cognition can be eliminated through short-term instruction or long-term experience in life (35-37). A longer duration of education resulted in increased cortical thickness in bilateral temporal, parietal, medial-frontal, sensory, and motor cortices in the adult brains (38). In addition, after six weeks of multi-component cognitive control training, adolescents showed increased gray matter volume in the right inferior frontal cortex (rlFC) (39). According to recent neuroimaging research, the frontoparietal region underpins WCST performance (40), and the frontal cortex is involved in attention, working memory, and set-shifting within this network (40, 41). Besides that, a previous study revealed that females benefit more from certain training than males (37), leading support to our findings in terms of brain morphology: the formal education which contains a curriculum for knowledge acquisition as well as physical and awareness development (42-44), boosts brain development in both sexes, particularly in the frontal cortex (45), and such effects are accentuated in women, attenuating sex differences in WCST test performances.

Functionally, we discovered that cholinergic signaling may be at the root of the effect of education on attenuating cognitive sex differences. Acetylcholine (ACh) is thought to be a neuromodulator in the central nervous system in adults, altering neuronal excitability, inducing synaptic plasticity, and coordinating the firing of groups of neurons (46). For example, the firing of dopamine (DA) neurons located in the ventral tegmental area, as well as the release of DA by their projections in the striatal area, is tightly controlled by the cholinergic pathway mediated by nicotinic acetylcholine receptors (nAChRs) (47). DA depletion has been linked to widespread impairment in connectivity between frontal cortex and striatum as well as worsened set-shifting ability during WCST (48), and this ability in WCST indicates the metric of learning, abstract reasoning, and problem solving, participants with better set-shifting have a better opportunity of achieving a higher correct rate (49). Furthermore, earlier research found that experience alters the cholinergic pathway in the hippocampus of mice by interaction with a gene product and that repeated learning specifically amplified the effect of such gene product expression on this cholinergic projection pathway (50). Effects of cholinergic enhancement on experience-dependent plasticity have also been observed in healthy adult auditory cortex (51, 52), as well as their visual perceptual learning (53), and a subsequent follow-up study elucidated that pharmacological cholinergic enhancement on visual perceptual learning is long-lasting (54). Baskerville and collaborators (55), on the other hand, investigated the effects of ACh input depletion from nucleus basalis on experience-dependent plasticity in the cortex of young adult male rats and discovered that cholinergic-depleted animals showed no significant plasticity response. This suggested that the cholinergic enhancement on learning-dependent plasticity is likely to be sex-specific, with females benefiting more.

Another finding of the present study is that nearly half of the DPEs interacted directly with estrogen. Not surprisingly, the cholinergic pathways are critical sites for estrogen in the brain (56), and multiple basic and preclinical research over the past decades has clearly demonstrated that basal forebrain cholinergic systems are relied upon estradiol support for adequate functioning (57), and these cholinergic projections play an essential role in attentional processes and learning (58, 59). Besides this, available evidence suggests many of the effects of estrogen on neuronal function and plasticity, as well as cognitive performance, are related to or dependent on interactions with such cholinergic projections (60-63). Consistent with these findings, we detected an estrogen-nAChR3 interaction that positively regulates cognitive performance, nAChR3 expression was upregulated in females and this upregulation was more remarkably in the secondary and tertiary education group.

Although the cholinergic-estrogen interaction offers the strongest evidence for the effect of education on attenuating cognitive sex differences. Other inherent or external factors with similar effects cannot be ignored. Prior research indicated that in rat, the arginine vasopressin (AVP) deficiency caused by a mutation in the *Avp* gene eliminated sex differences in the extinction of a conditioned taste aversion (64) and social reinforcement (65). Except for education, one sociocognitive factor was also proposed to abolish sex differences in mental rotation performance (66), the probable underlying mechanism might be confidence as a mediator by which sex stereotype exerts its influence, that is when females are the target of sex stereotype, their working memory capacity decreased dramatically (67, 68), resulting in a drop in logical reasoning (69) and mathematical performance (70).

There are some limitations to our study. First, even though this study included subjects ranging from young adults to elderly, the proportion of middle-aged and elderly subjects was much higher than that of young adults, furthermore, the young participants were clustered in the higher education group. A further study based on young adults is needed to replicate the current findings. Second, cognitive function and education are related, but not only because education influences cognition; higher intellectual function can lead to more education. Third, in this study, neuronal acetylcholine receptor subunit alpha-3 (nAChR3) was detected in the serum of the research subjects; however, due to the blood-brain barrier, the majority of the nAChRs may be synthesized locally; therefore, further research into the change of nAChRs expression levels in brain tissues induced by learning is required to replicate this study. Last, owing to the moderate sample size, further generalizing the results of our study need to be cautious.

## 5. Conclusion

This study investigated whether learning experience is mediated by sex-dependent neural activity. We discovered that education modifies the cholinergic signaling differently in males and females, with women benefiting more. Once again, this implies that brain development is the consequence of genetic-environmental interaction, and environmental factors can be a novel predictor for predicting and preventing specific mental disorders.

## Acknowledgements

CC was supported by the Naresuan Competitive Grants for International Students (NCG). We would like to thank Professor Gavin P. Reynolds, Biomolecular Sciences Research Centre, Sheffield Hallam University, UK, for his guidance and suggestions throughout the manuscript. We would also like to thank the facilities support from the Faculty of Medical Science, Naresuan University and The National Center for Genetic Engineering and Biotechnology, Pathum Thani, Thailand.

